# Widespread paleopolyploidy, gene tree conflict, and recalcitrant relationships among the carnivorous Caryophyllales

**DOI:** 10.1101/115741

**Authors:** Joseph F. Walker, Ya Yang, Michael J. Moore, Jessica Mikenas, Alfonso Timoneda, Samuel F. Brockington, Stephen A. Smith

## Abstract

- The carnivorous members of the large, hyperdiverse Caryophyllales (e.g. Venus flytrap, sundews and *Nepenthes* pitcher plants) represent perhaps the oldest and most diverse lineage of carnivorous plants. However, despite numerous studies seeking to elucidate their evolutionary relationships, the early-diverging relationships remain unresolved.
- To explore the utility of phylogenomic data sets for resolving relationships among the carnivorous Caryophyllales, we sequenced ten transcriptomes, including all the carnivorous genera except those in the rare West African liana family (Dioncophyllaceae). We used a variety of methods to infer the species tree, examine gene tree conflict and infer paleopolyploidy events.
- Phylogenomic analyses support the monophyly of the carnivorous Caryophyllales, with an origin of 68-83 mya. In contrast to previous analyses recover the remaining non-core Caryophyllales as non-monophyletic, although there are multiple reasons this result may be spurious and node supporting this relationship contains a significant amount gene tree discordance. We present evidence that the clade contains at least seven independent paleopolyploidy events, previously debated nodes from the literature have high levels of gene tree conflict, and taxon sampling influences topology even in a phylogenomic data set.
- Our data demonstrate the importance of carefully considering gene tree conflict and taxon sampling in phylogenomic analyses. Moreover, they provide a remarkable example of the propensity for paleopolyploidy in angiosperms, with at least seven such events in a clade of less than 2500 species.

## INTRODUCTION

Carnivory in plants has long fascinated both the general public and evolutionary biologists. Charles Darwin himself dedicated an entire volume to carnivorous species in his *Insectivorous Plants* (Darwin, 1875). The wide array of traps that are used to catch insects and other prey items make carnivorous plants some of the most morphologically diverse plants on Earth (Ellison and Gotelli, 2001; Heubl et al., 2006). These plants are able to occupy nutrient poor soils that would otherwise be unsuitable for plant life by obtaining nutrients unavailable in the soil through the digestion of animals.

Across angiosperms, carnivory is hypothesized to have independently evolved at least nine times (Givnish, 2015). One of these events is thought to have occurred relatively early on (~83mya) in the non-core Caryophyllales (Magallón et al., 2015), giving rise to a “carnivorous clade” consisting of the fully carnivorous families Droseraceae, Drosophyllaceae, and Nepenthaceae, the small non-carnivorous African family Ancistrocladaceae, and the rare west African family Dioncophyllaceae, which includes the unusual carnivorous liana *Triphyophyllum peltatum* and two other monotypic, non-carnivorous genera (*Dioncophyllum* and *Habropetalum*) (Albert et al., 1992; Meimberg et al., 2000; Brockington et al., 2009; Soltis et al., 2011; HernÁndez-Ledesma et al., 2015). This clade comprises approximately 250 of the estimated 600 species of carnivorous angiosperms (Heubl et al., 2006; Ellison and Gotelli, 2009) and includes a diverse assemblage of trap-plants and pitcher plants that occupy a wide range of ecosystems, from the fully aquatic *Aldrovanda vesiculosa* to desert species of *Drosera* to the rainforest liana *Triphyophyllum.* Moreover, carnivory also appears to have been lost 1-3 times (Heubl et al., 2006) within the carnivorous clade, including in the ancestor of the 16 species of Ancistrocladaceae (Taylor et al., 2005) as well as in the ancestors of *Dioncophyllum* and *Habropetalum* (Meimberg et al., 2000).

Despite broad appeal and interest, the evolutionary relationships in the non-core Caryophyllales remain ambiguous, with studies seeking to resolve these relationships often resulting in individually well supported but mutually conflicting topologies (Meimberg et al., 2000; Cameron et al., 2002; Brockington et al., 2009; HernÁndez-Ledesma et al., 2015). Much of this conflict involves the earliest branch in the non-core carnivorous clade, with studies finding Nepenthaceae as sister to the remaining lineages (HernÁndez-Ledesma et al., 2015), others finding Droseraceae as sister to the rest of the group (Meimberg et al., 2000), and yet others finding Droseraceae to be sister to the Nepenthaceae (Brockington et al., 2009). The strong support for conflicting topologies from different studies may be explained by the reliance on one or a few genes leading to systematic error (Maddison, 1997; Rokas et al., 2003). This type of error can arise from a variety of sources, including, but not limited to, incomplete lineage sorting, horizontal gene transfer, hybridization and hidden paralogy (Galtier and Daubin, 2008). Untangling these processes has proven to be a challenge and adds a strong level of complexity to phylogenomic analyses (Smith et al., 2015).

Transcriptomes have proven to be a powerful source of data for understanding this complexity, and have helped provide insight into the evolutionary history of non-model species (Dunn et al., 2008; Cannon et al., 2015; Yang et al., 2015). The thousands of genes typically sequenced in a transcriptome provide a means of identifying gene duplications and paleopolyploidy events (Cannon et al., 2015; Yang et al., 2015; Barker et al., 2016), which may clarify whether such events have been major drivers of evolutionary novelty (Ohno et al., 1968; Soltis et al., 2014). Moreover, analyses of gene tree concordance and conflict allows for a better understanding of the formation of species relationships and the complexity that arises in genomes as a result of speciation (Pease et al., 2016).

In this study, we conduct the first phylogenomic analysis focused on the non-core Caryophyllales, with sampling that covers all genera of carnivorous Caryophyllales except the poorly studied and rare liana *Triphyophyllum* (Dioncophyllaceae) of West Africa. We use large datasets to help resolve evolutionary relationships and explore gene tree discordance and its possible causes, as well as its consequences for phylogenetics among the carnivorous Caryophyllales. We find that, even with phylotranscriptomic data, many of the complications observed earlier in targeted sequencing studies (e.g. taxon sampling, gene tree conflict) are still present. However, we show how transcriptome data provide important insights into the reasons for these complications. Furthermore, we use transcriptome data to help provide information on the prevalence of polyploidy in this ecologically and morphologically diverse clade and explore the molecular evolution of the group.

## MATERIALS AND METHODS

### Data Availability

Raw reads for the ten newly generated transcriptomes were deposited in the NCBI Sequence Read Archive (Table S1; Bioproject: PRJNA350559). Assembled sequences, data files, programs, alignments and trees are available from Dryad (XXXX).

### Taxon Sampling, Tissue Collection, Sequencing and Data Assembly

The workflow for processing the data was run using a previously developed phylogenomic workflow (Yang et al., 2016). Transcriptomes of eight non-core Caryophyllales families representing nearly all of the major lineages of non-core Caryophyllales were included in this study (Table S1). The transcriptomes of *Dionaea muscipula, Aldrovanda vesiculosa, Nepenthes ampullaria* and *Reaumuria trigyna* were downloaded from the NCBI Sequence Read Archive [accessions SRX1376794, SRR1979677, (SRR2666506, SRR2866512 and SRR2866533 combined) and (SRX105466 & SRX099851 combined) respectively] (Dang et al., 2013; Brockington et al., 2015; Bemm et al., 2016; Wan Zakaria et al., 2016). The assembly used for *Frankenia laevis* was the same as in Yang et al. (2015) and can be found in Dryad (http://dx.doi.org/10.5061/dryad.33m48). The genomes of *Beta vulgaris* (RefBeet-1.2) and *Spinacia oleracea* were downloaded from The *Beta vulgaris* Resource (http://bvseq.molgen.mpg.de/Genome/Download/index.shtml; accessed Jul 10, 2015) (Dohm et al., 2014). We generated ten new transcriptomes for this study from fresh leaf tissue collected from *Drosera binata, Nepenthes alata, Ancistrocladus robertsoniorum, Plumbago auriculata, Ruprechtia salicifolia* and *Drosophyllum lusitanicum.* The *D. binata* and *N. alata* data were also collected from trap tissue at three different life stages. The plant tissues were flash frozen in liquid nitrogen and stored at −80°C. RNAs were extracted from the leaf tissue using the Ambion PureLink Plant RNA Reagent (ThermoFisher Scientific Inc, Waltham, Massachusetts, United States) following the manufacturer’s instructions and quantified using the Agilent 2100 Bioanalyzer (Agilent Technologies, Santa Clara, California, United States). Sequence libraries were prepared using the KAPA stranded mRNA library preparation kit (Kapa Biosystems, Wilmington, Massachusetts, United States) using the default protocols except for fragmentation at 94°C for 6 min and ten cycles of PCR enrichment. All ten libraries were multiplexed, then *D. binata* and *N. alata* were sequenced together on the same lane of the Illumina HiSeq2000 platform. *Ruprechtia salicifolia* was run on a separate Illumina HiSeq2000 lane with six other samples, *A. robertsoniorum* was run on a separate Illumina HiSeq2500V4 along with ten other samples and *P. auriculata* was run on a separate Illumina HiSeq2500V4 run along with ten other samples.

The raw paired end reads from the newly generated transcriptomes were trimmed and filtered using Trimmomatic (Bolger et al., 2014) with trim settings sliding window 4:5, leading 5, trailing 5 and min length 25. For both *D. binata* and *N. alata,* the three transcriptomes from trap tissues were combined and assembled together. The procedure was conducted as follows: the remaining read set was assembled using Trinity v2.04 (Grabherr et al., 2011) with strand-specific settings and stranded ‘RF’ and the assembled reads were translated using Transdecoder v2.0 (Haas et al., 2013) guided by BLASTP against a BLAST database consisting of concatenated *Arabidopsis thaliana* and *B. vulgaris* proteome (Dohm et al., 2014), with strand-specific settings. All translated amino acid datasets were reduced with cd-hit v4.6 (-c 0.995-n 5) (Fu et al., 2012).

### Analysis of Sources of Contamination

We tested for within-lane contamination by creating one-to-one ortholog gene trees (using the pipeline described below) and comparing the resulting tree topologies to the expected species tree topology for all samples on the lane. Additionally, we examined *matK* sequences from the assembled transcriptome coding DNA sequence (CDS) data. Using these sequences together with those obtained from GenBank (Table S2) to represent each of the non-core families used in the analysis, we constructed a phylogeny using maximum likelihood and the settings “-f a -# 200-m GTRCAT-p 12345-x 112233” as implemented in RAxML (Stamatakis, 2014). We were unable to recover *matK* from two of the assembled transcriptomes (*A. vesiculosa* and *P. auriculata*), and instead we ensured that the highest GenBank BLAST hit was that of the same species *A. vesiculosa* (AY096106.1) and *P. auriculata* (EU002283.1) respectively.

### Homology Inference and Species Tree Estimation

Homology and orthology inference along with species tree estimation were carried out following Yang and Smith (2014), which is briefly summarized below. The exact commands and programs are available either at https://github.com/jfwalker/JFWNonCoreCaryophyllales for scripts involved in the downstream analysis or at https://bitbucket.org/yangya/phylogenomic_dataset_construction for scripts used in assembling the species tree. After the peptide and coding DNA sequences were reduced using cd-hit, we created six datasets to explore the influence of taxon sampling and sequence type. Three of the datasets were made using the peptide data. One dataset consisted of all taxa; one dataset excluded *Ancistrocladus robertsoniorum* and one dataset excluded *Drosophyllum lusitanicum.* We then created corresponding nucleotide sequence datasets with the same taxon content. All steps for the homology inference and species tree estimation were the same for all datasets, except where noted below. The first step was an all-by-all BLASTP search, in the case of the peptide datasets, or an all-by-all BLASTN search in the case of the nucleotide data, which was conducted with an e-value of 10. Putative homolog groups were formed by retaining species with a hit fraction >0.4 and using Markov clustering as implemented in MCL14-137 (Van Dongen, 2000) with the inflation value set to 1.4 and e-value cutoff of 5. Only clusters that had at least 4 taxa were retained.

Each cluster was then aligned using MAFFT v7 (Katoh and Standley, 2013) with “-genafpair maxiterate 1000” and trimming of the alignments was conducted using Phyutility v2.2.6 (Smith and Dunn, 2008) with “-clean 0.1”. For sequence clusters containing less than 2000 sequences, the phylogenetic trees were estimated through maximum likelihood as implemented in RAxML v8.2.3 (Stamatakis, 2014) with the model PROTCATWAG (AA) or GTRCAT (DNA). In the case of sequence clusters larger than 2000 sequences, this was done with FastTree 2 (2.1.8) (Price et al., 2010) with the WAG model (AA) or the GTR model (DNA). All single branches greater than 2 substitutions per site were removed as these are likely the result of sequences being pulled together by error or conserved domains. We also removed all branches 10 times or greater in length than their sister branches in the homolog tree for similar reasons. In the case of clades, the analysis took the step-wise average from root to tip and removed it if that was greater than 10 times the length of the sister and the tips of the same species that appeared as monophyletic, indicating they were likely alternate transcripts or in-paralogs. Further data refinement was done by removing all the monophyletic tips except the tip associated with the sequence with the highest number of aligned characters after trimming (i.e. most informative) data. The sequence data were then removed from the homolog trees and the process was repeated a second time, to further clean the data.

The support for the homolog trees was analyzed after the second round using the Shimodaira-Hasegawa-like approximate likelihood ratio branch test (Anisimova et al., 2011) as implemented in RAxML, for downstream analysis only branches with (SH-Like => 80) we considered informative. Then one-to-one orthologs were identified from the homolog trees (Yang and Smith, 2014), using *B. vulgaris* and *S. oleracea* as outgroups, both of which are in the core Caryophyllales and have genome information. The ortholog trees produced from these methods were then used to extract the amino acid sequence data associated with the given ortholog tree. A dataset was created from one-to-one orthologs containing no missing taxa. Each ortholog produced from each method was then individually aligned using PRANK v. 140603 with default parameters (Löytynoja and Goldman, 2008). The alignments were then trimmed using Phyutility with a minimum occupancy of 0.3 being required at each site. Supermatrices were created for all approaches by concatenating all trimmed alignments that had at least 150 characters. A maximum likelihood tree for each supermatrix was estimated using RAxML with the PROTCATWAG model, partitioning by each ortholog group. Node support was evaluated using 200 nonparametric bootstrap replicates. Following this the Maximum Quartet Support Species Tree (MQSST) was found using ASTRAL (v4.10.0) (Mirarab et al., 2014) with default parameters and using the one-to-one ortholog trees as the inputs.

### Dating Analysis

To conduct the analysis, we used the 1237 orthologs identified in the nucleotide dataset and first found the genes whose gene tree matched the species tree. From the 135 genes that met this criterion, we calculated the variance from each tip to root, using pxlstr from the Phyx package (Brown et al. *in review*). The dating analysis was conducted using BEAST (ver. 1.8.3) (Drummond and Rambaut, 2007) on the three genes with the lowest variance as they represent the genes evolving in the most clocklike manner. We used the GTR+G model of evolution and a birth-death tree prior. We calibrated the clade containing the genera Aldrovanda and Dionaea with a lognormal prior with offset 34 and a mean of 0 and standard deviation of 1 based on a fossil Aldrovanda (Degreef, 1997). Because of the low root to tip variance for the three genes (~0.0004), we used the strict clock model for the rates of evolution. We ran the MCMC for 10,000,000 generations and the first 1,000,000 generations were discarded as the burn-in. We summarized the topology as the maximum clade credibility tree.

### Gene Family Size Analysis

Two sets of gene families were analyzed, one for the overall largest gene family and one for the gene families previously associated with the adaptation to carnivory in a differential gene expression study (Bemm et al., 2016). To identify the overall largest family, we found the inferred homolog trees that had the largest number of tips, and annotation was done by taking a representative sample from the homolog tree and finding the highest hit on NCBI blast database. For the carnivorous gene families, representative samples from the genes identified in *Bemm et. al* were downloaded from Genbank (Table S3). A blast database was created from the downloaded samples and BLASTP was used to identify their corresponding sequences, which were then found in the homologous gene clusters. The number of tips were counted for each homologous gene tree to identify the size of the gene family and number of genes associated with carnivory.

### Analysis of Gene Duplications

Gene duplications were analyzed with phyparts (vrs. 0.0.1) (Smith et al., 2015) using the homolog clusters. Only gene duplications with nodes that contained (≥80) SH-Like support were used to identify duplications. The homolog clusters for each of the six datasets were mapped onto their respective species tree topologies. Further analysis of the gene duplications was conducted by finding all gene duplications, irrespective of species tree topology, using a modified version of phyparts. Again in this case only gene duplications that contained (≥80) SH-Like support were removed from the homolog trees. These duplications were then used to create a phylogenetic tree by creating a shared presence matrix from existing duplications and correcting for distance by taking (1/number of shared duplications). The distance matrix was used to create a phylogenetic tree following the Neighbor-Joining method (Saitou N, 1987). The modified version of phyparts and script (GeneJoin.pl) that creates a phylogenetic tree from that output can be found at (https://github.com/jfwalker/JFWNonCoreCaryophyllales).

### Analysis of Gene Tree Conflict

The one-to-one orthologs recovered from the homolog trees were used to analyze the gene tree/species tree conflict at all nodes and this analysis was performed on all six datasets, with their respective gene trees and species tree being used for each individual analysis. The orthologs were all rooted based on *S. oleracea* and *B. vulgaris* using the phyx program pxrr (Brown et al., 2017). The rooted one-to-one ortholog trees were then compared to the species tree using phyparts with only informative branches being counted. The output of phyparts was used to identify the amount of conflict at each node along with the dominant alternative topology.

### Inferring genome duplication events

To infer potential genome duplication events, we visualized the number of synonymous substitutions that were found between the paralogs with all of the taxa. The process was carried out using the script ks_plots.py from Yang et al. 2015 (https://bitbucket.org/yangya/carvophyllalesmbe2015) which relies upon the pipeline from (https://github.com/tanghaibao/bio-pipeline/tree/master/synonymouscalculation). The pipeline first reduces sets of highly similar sequences using CD-HIT (-c 0.99-n 5). Following this, an all-by-all BLASTP is carried out within each taxon using an e-value of 10 and-max_target_seq set to 20. The resulting hits with < 20% identity or niden < 50 amino acids are removed. The sequences that have ten or more hits are removed to avoid over representation of gene families. The remaining paralog pairs are then used to infer the genome duplications, as areas where the Ks value is greater than the background rate (Schlueter et al., 2004). First pairwise protein alignments are created using the default setting of ClustalW (Larkin et al., 2007), these are then back translated to codon alignments using PAL2NAL, and the synonymous substitutions rates are calculated using yn00 of the PAML package (Yang, 2007), with Nei-Gojobori correction for multiple substitutions (Nei and Gojobori, 1986).

To infer the phylogenetic locations of genome duplications, we used a comparison of the genome duplication events identified from paralogs mapped onto the Ks plots of multiple species made from the reciprocal blast hits. The process was carried out using the script MultiKs.pl, which can be found at (https://github.com/jfwalker/JFWNonCoreCaryophyllales). The pipeline works as follows. First the highly similar sequences are reduced using CD-HIT (-c 0.99-n 5). Then a reciprocal BLASTP is carried out on the peptide transcriptomes where one of the transcriptomes is used as a query and another is used as the database. Following that the top blast hit is removed and the peptide sequences are aligned using MAFFT. The peptide alignment is then matched with the corresponding nucleotide files and the nucleotides are aligned based on the peptide alignment using the phyx program pxaatocdn (Brown et al., 2017). From there the synonymous substitution rates are calculated using yn00 of the PAML package, with the Nei-Gojobori correction for multiple substitutions. The Ks peaks of the genome duplications inferred from the paralogs are then compared to the Ks peaks of the multispecies comparison, if the peak from the single species comparison is smaller than the multi-species, this provides evidence that the genome duplication occurred after the speciation event (Cannon et al., 2015).

### Comparing molecular rates among differing gene tree topologies

The gene trees that contained the topologies supporting either *Drosophyllum* and *Ancistrocladus* as sister to all other lineages or *Drosophyllum* and *Ancistrocladus* as sister to *Nepenthes* were identified from the bipartitions removed using the phyx program pxbp (Brown et al., 2017) and the program GeneHybridSplitter.pl (https://github.com/jfwalker/JFW_NonCoreCaryophyllales). The ortholog tree was considered to support *Drosophyllum* and *Ancistrocladus* as the lineage sister to the others if it contained a bipartition containing only *Drosophyllum* and *Ancistrocladus*, a bipartition containing only the carnivorous lineages except *Drosophyllum* and *Ancistrocladus*, and a bipartition containing only and all the carnivorous taxa. The ortholog trees that supported *Drosophyllum* and *Ancistrocladus* sister to *Nepenthes* were identified if the tree contained a bipartition with only *Ancistrocladus* and *Drosophyllum*, a bipartition with both *Nepenthes* species and *Drosophyllum* and *Ancistrocladus,* and a bipartition containing only and all the carnivorous taxa.

The synonymous substitution rates found in both scenarios were calculated using a pairwise comparison of *Drosophyllum* and *Nepenthes alata,* along with a pairwise comparison of *Ancistrocladus* and *N. alata.* The corresponding nucleotide and amino acid sequences of *Drosophyllum* and *N. alata* were removed for all the gene trees that support *Ancistrocladus* and *Drosophyllum* as the basal lineage. The pairwise amino acid sequences were then aligned using MAFFT, and the amino acid alignment was then used to guide the codon based alignment using pxaatocdn. The Ks values for each codon alignment were calculated using the script Ks_test.pl (https://github.com/jfwalker/JFW_NonCore_Caryophyllales), which uses yn00 from the PAML package to obtain the Nei-Gojobori correction for multiple substitutions Ks values. The same procedure for finding synonymous substitutions was then performed on pairwise comparisons of *Drosophyllum* and *N. alata*, where they appear as sister, and was performed on *Ancistrocladus* and *N. alata* for the same situations.

## RESULTS

### Species tree, dating analysis and gene tree conflict

The monophyly of the non-core Caryophyllales was supported in both the concatenated maximum likelihood supermatrix (Fig. S1) and the maximum quartet support species tree (MQSST) reconciliations (Fig. S2), regardless of taxon sampling or molecule type used in the analysis. The divergence of this group appears to have occurred ~90 mya ago, with adaptation of carnivory arising ~75 mya (Fig. 1). A general trend was that branches of high conflict resulted in shorter branch lengths for both the concatenated supermatrix and the MQSST analysis (Fig. S1, S2). A clade of Frankeniaceae and Tamaricaceae was supported as sister to the remaining non-core Caryophyllales in all datasets by most gene trees. In the case of the ALLTAX AA dataset, the branch supporting this as the lineage sister to everything else showed a large amount of conflict with ~15.4% of genes supporting the topology, ~14.6% supporting a dominate alternate topology of a monophyletic non-carnivorous non-core (NCNC), ~25% supporting other alternate topologies and ~45% of gene trees being poorly supported (SH-Like < 80), with similar results for the five other datasets used to reconstruct the species tree topology. Further support of a non-monophyletic relationship of the NCNC was obtained by looking at the number of uniquely shared gene duplications found by the AA ALLTAX for the families in the carnivorous non-core, which in the case of Plumbaginaceae and Polygonaceae was 103. This is in contrast to the five unique gene duplications shared among the NCNC. The MQSST and concatenated ML supermatrix analyses inferred that the next lineage to diverge was a clade containing both the families Plumbaginaceae and Polygonaceae, whose sister relationship received 100% bootstrap support and ~70% genes concordant with the topology with 10.5% conflicting in the case of the AA ALLTAX. This relationship showed up in all datasets regardless of composition of taxa used for the analysis.

**Figure 1.**
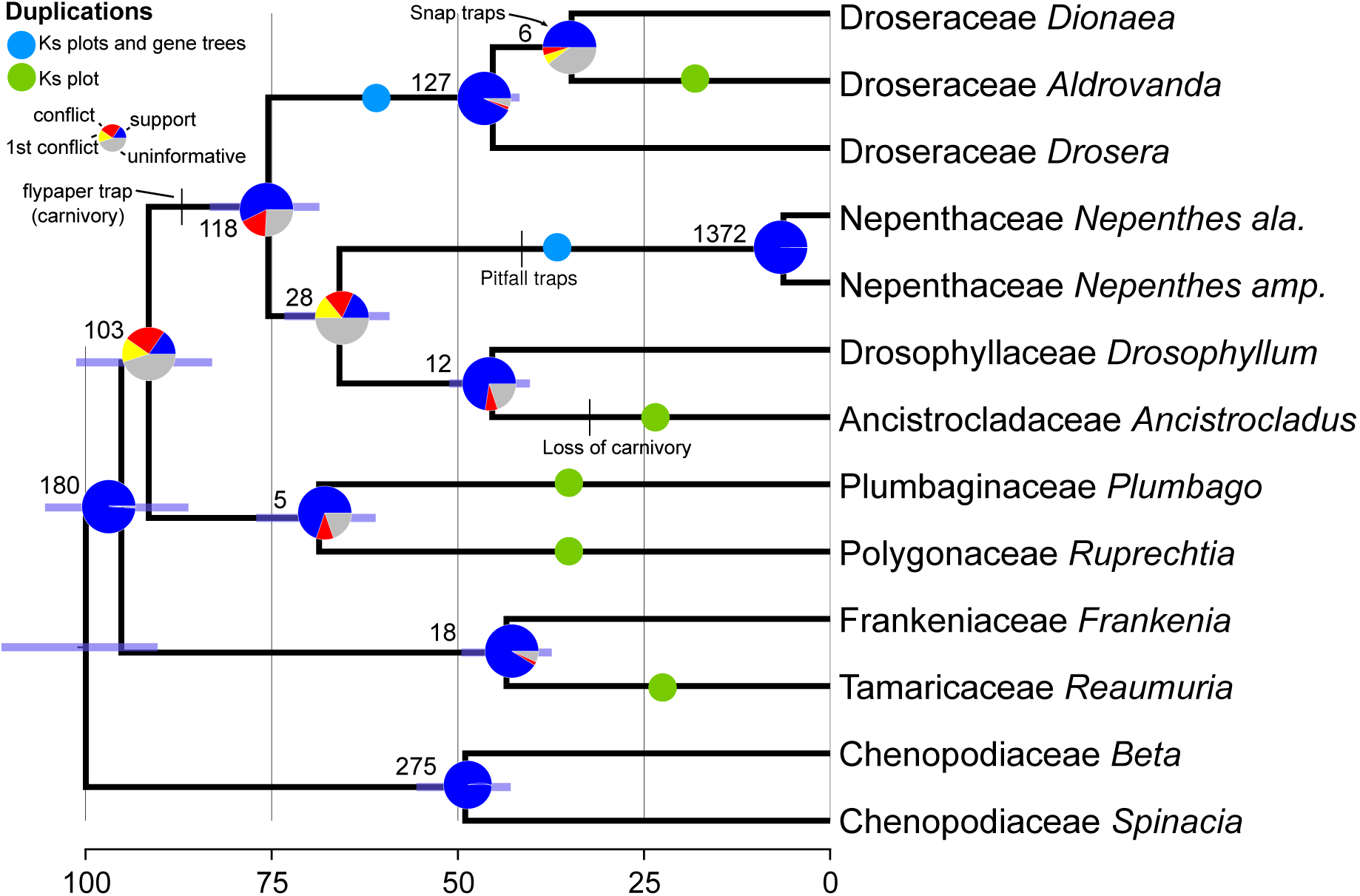
Inferred and dated species tree from the three-gene Bayesian dating analysis. Numbers on each branch represent inferred shared unique to clade gene duplications, and branch lengths are proportional to time. Circles on branches represent inferred genome duplications, position supported only by Ks plots (Green) and position supported by Ks plots along with shared gene duplications (Blue). Pie charts show gene tree conflict evaluations at each node, proportion concordant (Blue), proportion conflicting (Red), dominant alternative topology (Yellow) and unsupported with SH-Like less than 80 (Grey). Ancestral states on branches taken from *Heubl et. al 2006*.

All datasets revealed a strongly supported (BS = 100%) clade consisting of the carnivorous families and the non-carnivorous family Ancistrocladaceae. In the case of the AA ALLTAX dataset the majority of the well-supported gene trees (~57%) were concordant with the species tree topology, with similar results for all other datasets. In all cases, Droseraceae and Nepenthaceae were each monophyletic (Fig. 2). The main discordance in the species tree topology involved the placement of Drosophyllaceae (Fig. 2). When all taxa were included Drosophyllaceae was sister to Ancistrocladaceae, a relationship that is well supported by concordant gene signal in both the AA dataset (72.5%) and the CDS dataset (93.7%). However, the placement of the clade containing Drosophyllaceae and Ancistrocladaceae changed depending on sequence type: for AA data it is reconstructed as sister to the Nepenthaceae, whereas for CDS data it is sister to the rest of the carnivorous clade, albeit with no bootstrap support (Fig. 2).

**Figure 2.**
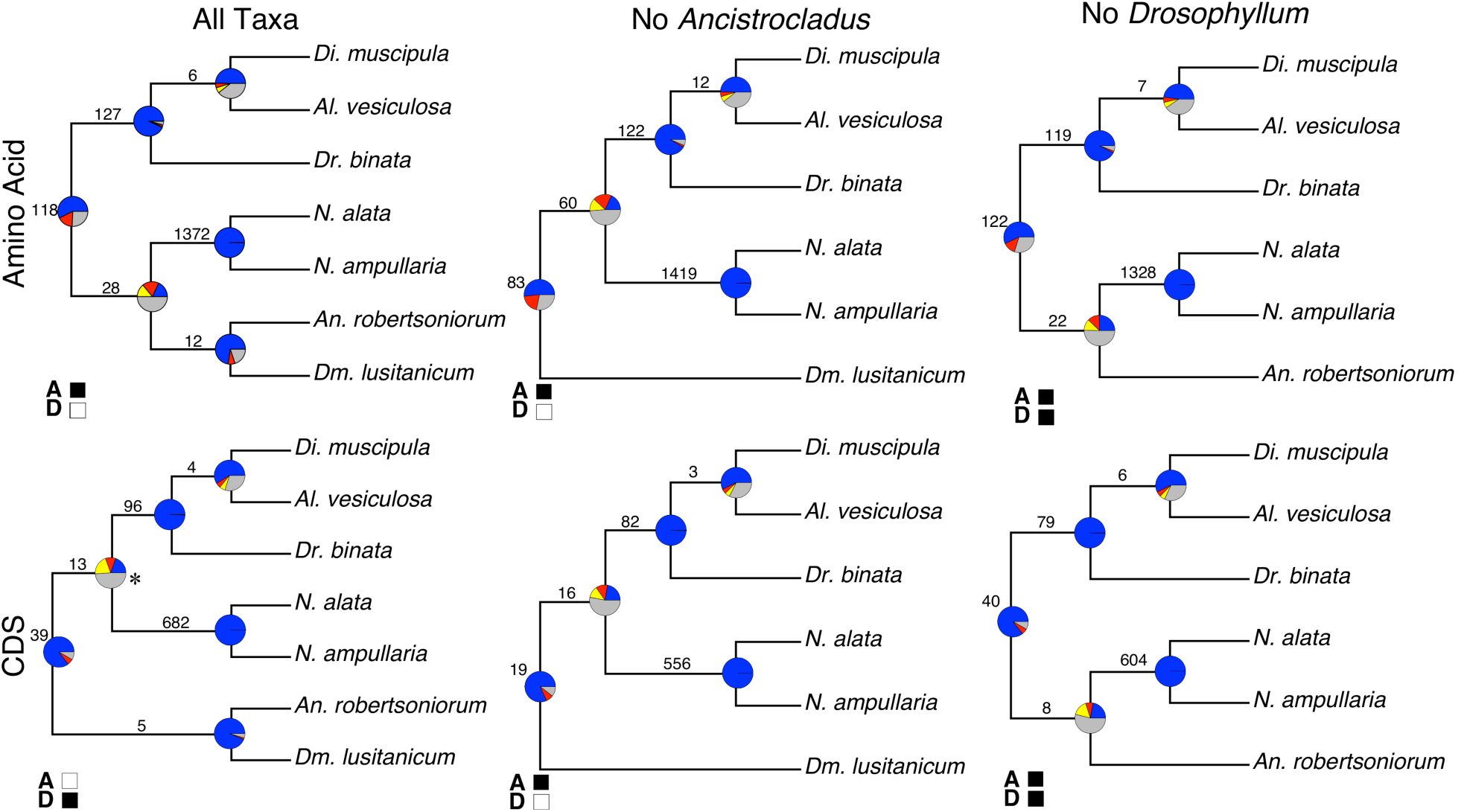
The influence of taxon sampling and sequence type on inferred tree topology. Respective topologies are from the RAxML supermatrix analysis, filled boxes are used to represent concordance with a different method of species tree reconciliation “A” represents Astral (MQSST) and “D” represents Distance matrix reconstruction. Star near the node indicates BS support of 0, all other nodes have BS support of 100. Numbers on each branch represent inferred gene duplications. Pie charts show gene tree conflict evaluations at each node, proportion concordant (Blue), proportion conflicting (Red), dominant alternative topology (Yellow) and unsupported with SH-Like less than 80 (Grey).

When *Ancistrocladus* was excluded from analyses, for both the AA and CDS datasets, Drosophyllaceae appeared as sister to the rest of the taxa in the carnivorous clade (Fig. 2b, e). The clade containing Droseraceae and Nepenthaceae has a large amount of discordance with ~18% concordant and 32% conflicting for the AA dataset and ~20% concordant and ~22% conflicting for the CDS dataset. In both cases this was a node where many of the gene trees contained low Shimodaira-Hasegawa-Like support (< 80%). When *Drosophyllum* was excluded from analyses, for both the CDS and the AA datasets, Ancistrocladaceae appeared as sister to Nepenthaceae. Again, the node that defined this relationship had a significant amount of conflict, where in the AA dataset ~25% of the gene trees showed a concordant topology and ~24% showed a conflicting topology. With the CDS dataset ~22% of gene trees were concordant with the species topology and ~24% gene trees were conflicting. Again in both cases many of the gene trees did not have strong SH-Like (≥80) support for either topology.

### Analysis of potential hybridization and comparison of synonymous substitutions rates (KS) between woody and herbaceous species

No differences were found between the synonymous substitution rate between the gene trees supporting the sister position of *Drosophillum lusitanicum* and *Aldrovanda robertsoniorum* to the remaining lineages as opposed to those supporting the two species as sister to only Nepenthaceae (Fig. S3). For *D. lusitanicum,* the mean Ks for the trees supporting the sister to the other lineages position was 0.8546, whereas those supporting the position sister to Nepenthaceae had a mean Ks value of 0.8586. In the case of *A. robertsoniorum* those supporting a sister to the other lineages relationship had a mean Ks value of 0.6359 and those supporting a relationship sister to only Nepenthaceae is 0.6358.

### Genome duplications and gene family sizes

The single-species Ks plots showed that all the Caryophyllales have at least one peak around 2.0 (Fig. S4). These plots also showed one additional peak for all taxa in non-core Caryophyllales except for *A. vesiculosa,* which had two additional peaks, and both *D. lusitanicum* and *Frankenia laevis* did not show any extra peaks. A comparison of Ks values between orthologs and paralogs for species pairs showed that in the case of Plumbaginaceae and Polygonaceae, the genome duplication likely occurred post speciation (Fig. 3). This post speciation genome duplication received further support as the two species only shared five unique gene duplications. This same comparison for representative species pairs of Ancistrocladaceae-Nepenthaceae and Droseraceae-Nepenthaceae showed that these genome duplications likely occurred after the divergence of the respective families in each pair (Fig. 3). An among Droseraceae comparison showed the duplication to have occurred after speciation in *Dionaea* but before speciation in *Drosera* (Fig. S5). The peak for the duplication appeared to be before-speciation in a comparison to *Drosera* and *Aldrovanda* (Fig. S5). Overall, the shared unique gene duplications and Ks plots support the inference of seven separate genome duplications across the non-core Caryophyllales, with six occurring after divergence of the families and none being uniquely shared by any two families in the group (Fig. 3).

**Figure 3.**
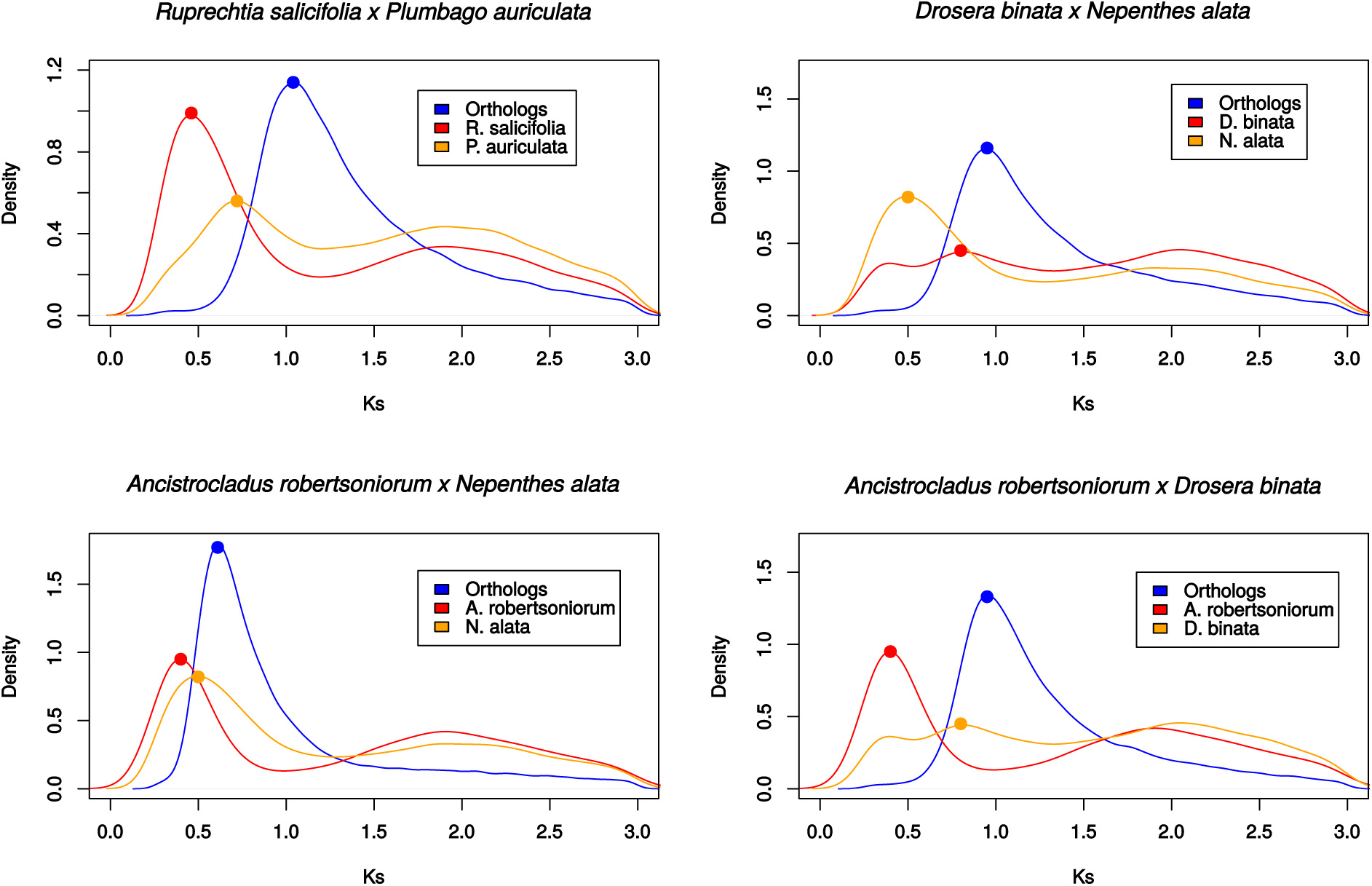
Representative Ks plots. Density plots representing the peak of the Ks values inferred from reciprocal orthologs (Blue) and those inferred from the within species paralogs (Red and Orange), with the density calculated for Ks values (=>0.25).

An analysis of the size of homologous gene families on the AA ALLTAX dataset showed that the largest gene family consisted of 3498 homologs (Table S4) and this family was associated with the function “putative leucine-rich repeat receptor-like protein kinase”. When further broken down into genes that are associated with carnivory, we found that the largest of these gene families was the “Plant Peroxidase” family (Table S5). On average, we did not find any specific gene family to have a disproportionate number of duplicated genes in the carnivorous plants as compared to the rest of the samples in the remaining non-core Caryophyllales, however, the plant peroxidase family has shrunk in the carnivorous lineage.

### Contamination checking and homology and orthology inference

Three major steps were taken to ensure that we would minimize the possibility of contamination in our samples. The first step was to extract the RNAs, prepare the sequencing libraries, and sequence the samples on separate lanes at different times. This was done for all samples we processed in this study other than *Nepenthes alata, Drosera binata,* and the previously published *D. lusitanicum,* which were sequenced together on a single lane. The next step was to create one-to-one ortholog phylogenetic trees out of the samples that were on the same lane, which showed most gene trees support previously accepted hypotheses for the often distantly related species on the lane. The final step was to ensure that the *matK* sequence from each of our assembled transcriptome shared the closest evolutionary relationship with a *matK* sequence taken from the same genus for each sample (Fig. S6; Table S2).

The datasets were made of the following taxon compositions for both amino acid (AA) and coding DNA sequence (CDS): all 13 taxa included (ALLTAX), all taxa except *D. lusitanicum* (NO DROS), and all taxa except *A. robertsoniorum* (NO ANC). The two datasets with all 13 taxa revealed that the inferred number of homolog clusters containing at least four taxa was the greatest using nucleotide data (Table S6). This is in contrast with both datasets that consisted of 12 taxa, in which the amino acid datasets inferred more homolog clusters than the nucleotide datasets. The complete taxa one-to-one orthology inference was comparable between all datasets of different taxa composition, where each time the amino acid dataset detected roughly 400 more one-to-one orthologs than its corresponding nucleotide dataset (Table S6).

## DISCUSSION

### Discordance among species trees and gene trees

Our transcriptome data confirm the monophyly of the carnivorous clade of Caryophyllales detected in previous studies (Meimberg et al., 2000; Brockington et al., 2009) and imply an ancient origin for the group, which our analyses suggest originated between 68-83 mya (Fig. 1). Our analyses further confirm that carnivory was the likely ancestral character state for the carnivorous clade, and that a mucilage trap characterized the progenitor of this clade (Heubl et al., 2006). Nevertheless, the subsequent evolution of life history within the carnivorous clade is less certain because it depends upon the topology of the earliest branches within the group, which have been unstable in previous analyses (Meimberg et al., 2000; Brockington et al., 2009; HernÁndez-Ledesma et al., 2015).

The large datasets generated in our study provide unique insight into the sources of this topological instability (Galtier and Daubin, 2008). For example, the shifting phylogenetic placement of *D. lusitanicum* could result from events such as horizontal gene transfer, incomplete lineage sorting, and/or ancient hybridization between an ancestral lineage that diverged prior to the other carnivorous Caryophyllales and one that diverged after the speciation event between Ancistrocladaceae and Nepenthaceae. The Nepenthaceae provides a logical source of hybridization as many of the species in genus are still capable of producing viable hybrids and do so in the wild (McPherson, 2009). If hybridization were the cause, we would expect two points of coalescence between *D. lusitanicum* and *N. alata* that would be associated with different synonymous substitution (Ks) values, as they would be influenced by the amount of time there was shared common ancestry with *N. alata.* An examination of Ks values did not reveal a difference in Ks values between the gene trees supporting the sister to all other lineages position or the sister to only Nepenthaceae position from the nucleotide data for either *D. lusitanicum* or *A. robertsoniorum* (Fig. S3). This provides some evidence that something other than hybridization may be the cause. However, full genome sequences would be necessary to improve confidence in our ability to discriminate among these processes because they would allow for direct association of phylogenetic signal over contiguous regions of chromosomal space (Fontaine et al., 2015). However, we did find that Ks values varied greatly between the *D. lusitanicum* and *A. robertsoniorum* comparisons, which may result from differences in habit, with the lineage of *Ancistrocladus* + Dioncophyllaceae transitioning to lianas and *Drosophyllum* retaining the ancestral herbaceous life history (Smith and Donoghue, 2008; Yang et al., 2015).

The remaining families of non-core Caryophyllales (Polygonaceae, Plumbaginaceae, Tamaricaceae, and Frankeniaceae) have previously been inferred to be a clade (Meimberg et al., 2000; Brockington et al., 2009; Soltis et al., 2011; HernÁndez-Ledesma et al., 2015; Yang et al., 2015), but our transcriptome-based analyses suggest that the clade of Frankeniaceae and Tamaricaceae and that of Plumbaginaceae and Polygonaceae are successively sister to the carnivorous clade. It is possible that this conflict is the result of our study including more informative phylogenetics characters in the analysis. However, it may also be the result of our relatively limited taxon sampling for these families and/or from the large number of conflicting gene trees associated with divergence events among these three groups (Fig. 1). The large number of conflicting gene trees may, itself, be the result of ILS associated with the relatively rapid divergence of these groups, as demonstrated by the short branch lengths from the MQSST analysis and concatenated supermatrix analysis (Fig. S1, S2). The uniquely shared gene duplications provide evidence for the sister relationship between the carnivorous clade and the clade of Plumbaginaceae + Polygonaceae. However, it should be taken into account that the higher number of gene duplications shared between Plumbaginaceae, Polygonaceae and the carnivorous Caryophyllales could be the result of biased sampling from more thorough sequencing, as transcriptomes are typically only found to recover up to half of coding genes (Yang and Smith, 2013). This provides a potentially biased sample for data when looking at uniquely shared gene duplications.

The disagreement between the supermatrix and MQSST methods of species tree reconciliation was likely a product of how the genes were treated in the analyses. In the MQSST all genes are given equal weight regardless of their informativeness and strength of the phylogenetic signal provided by the characters that created them, whereas in the supermatrix approach more informative genes provide a stronger signal for the overall matrix. The conflicting node for the CDS topology, however, received no bootstrap support.

Our results help to illustrate the important role that taxon sampling plays even when using character-rich datasets such as those used in phylogenomic reconstructions. In the analyses presented here, *D. lusitanicum* changed positions depending on the sampling used (Fig. 2). This discrepancy was not identified by the non-parametric bootstrap method, as 100% support was given to all nodes in all the reconstructions using the amino acid datasets, regardless of the position of *D. lusitanicum*. This helps to emphasize the importance of looking at more than just the non-parametric bootstrap in phylogenomic reconstructions, as in our datasets it is prone to Type I error and using transcriptome data allows us to examine conflicting signals. The non-parametric bootstrap, however, provided no support for the conflicting signal produced from nucleotide data. While we are unable to include Dioncophyllaceae in our analyses because of the difficulty in obtaining tissue, it is unlikely that inclusion would dramatically change carnivorous relationships given the strong support for its sister relationship to Ancistrocladaceae in all previous analyses (Heubl et al., 2006; Brockington et al., 2009).

### At least seven independent paleopolyploidy events in a group of less than 2500 species

Over the past decade, ever-larger phylogenomic datasets and improved methods for detecting genome duplications have revealed that paleopolyploidy is much more common in plants than previously thought (Barker et al., 2008, 2016; Yang et al., 2015). Previous evidence has suggested that the non-core Caryophyllales contain at least three paleopolyploidy events (Yang et al., 2015). Genome duplications have previously been implicated to be a source of novelty (Freeling and Thomas, 2006; Edger et al., 2015), a source of increased diversification (Tank et al., 2015), and decreased diversification (Mayrose et al., 2011). The seven inferred genome duplications of our analysis indicate that genome duplication has been a common occurrence in the history of the non-core Caryophyllales and is especially prevalent considering the group is estimated to have less than 2500 species (Soltis et al., 2006). Our results also support a shared genome duplication between the core and non-core Caryophyllales giving support to the evidence that at least one duplication occurred at the base of the group (Dohm et al., 2012). From our dataset it appears most of the non-core Caryophyllales families have unique genome duplication events. We found a discrepancy in the location of the duplication when comparing *Drosera* to *Dionaea* and when comparing *Drosera* to *Aldrovanda.* This may be due to the duplication occurring shortly before speciation or to the difference in rates of evolution found between *Aldrovanda* and *Dionaea* (Fig. S1). Without exhaustive sampling of each family it will not be possible to pinpoint the phylogenetic locations of the putative duplication events and hence it is not currently possible to determine whether a given paleopolyploid event acted to drive speciation and/or promote ecophysiological and morphological novelty. Nevertheless, the rich diversity and large number of genome duplications present within the non-core Caryophyllales suggests that this group will be a powerful tool for understanding genome and phenome evolution.

## ACKNOWLEDGEMENTS

We thank Edwige Mayroud, Joseph Brown, and Oscar Vargas for thoughtful comments on the manuscript and Ning Wang, Sonia Ahluwalia, Jordan Shore, Lijun Zhao, Alex Taylor and Drew Larson for helpful discussion on the manuscript; M. Raquel Marchán Rivadeneira for help with lab work; and Deborah Lalumondier and Justin Lee at the Missouri Botanical Garden for access to their living collections. The molecular work of this study was conducted in the Genomic Diversity Laboratory of the Department of Ecology and Evolutionary Biology, University of Michigan. This work was supported by NSF DEB awards 1352907 and 1354048.

## AUTHOR CONTRIBUTIONS

J.F.W., Y.Y., M.J.M., S.F.B. and S.A.S designed research. Y.Y., S.F.B., M.J.M. contributed to sampling; Y.Y., A.T. and J.M. conducted lab work; J.F.W. and Y.Y. performed sequence processing; J.F.W. and S.A.S. analyzed the data and led the writing.

## ONLINE SUPPLEMENTAL MATERIAL

**Supplementary Figure 1:**
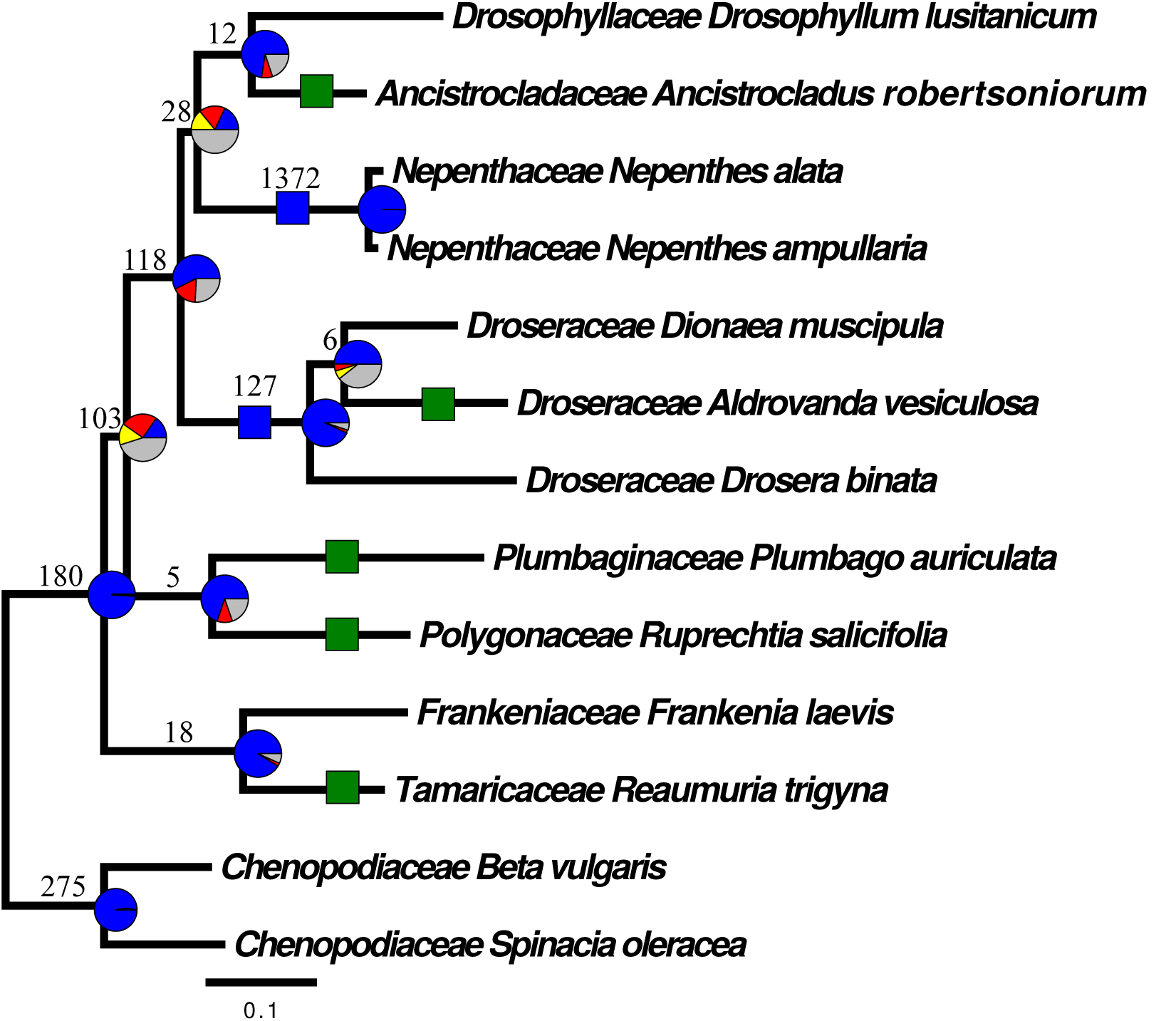
Species tree from RAxML analysis of the ALLTAX AA supermatrix. Numbers on each branch represent inferred shared unique to clade gene duplications. Squares along branches represent inferred genome duplications, position supported only by Ks plots (Green) and position supported by Ks plots along with shared gene duplications (Blue). Pie charts show gene tree conflict evaluations at each node, proportion concordant (Blue), proportion conflicting (Red), dominant alternative topology (Yellow) and unsupported with SH-Like less than 80 (Grey). Ancestral states on branches taken from *Heubl et. al 2006.*

**Supplementary Figure 2:**
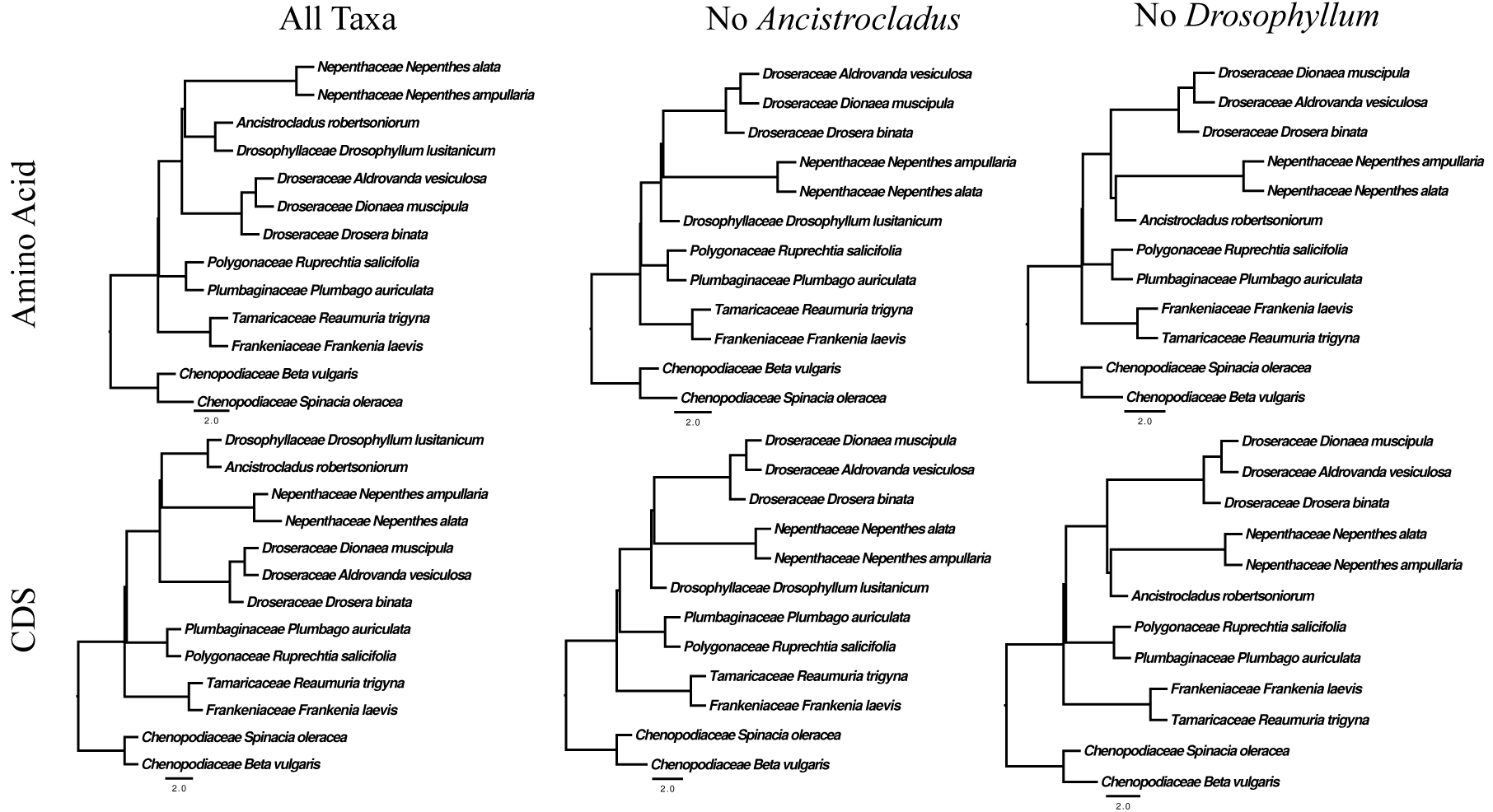
Inferred species trees from the Maximum Quartet Supported Species Tree analyses as implemented in Astral. The figure shows the different topologies that result from different combinations of molecules and species sampling inferred using the Maximum Quartet Supported Species Tree (MQSST) as implemented in Astral.

**Supplementary Figure 3:**
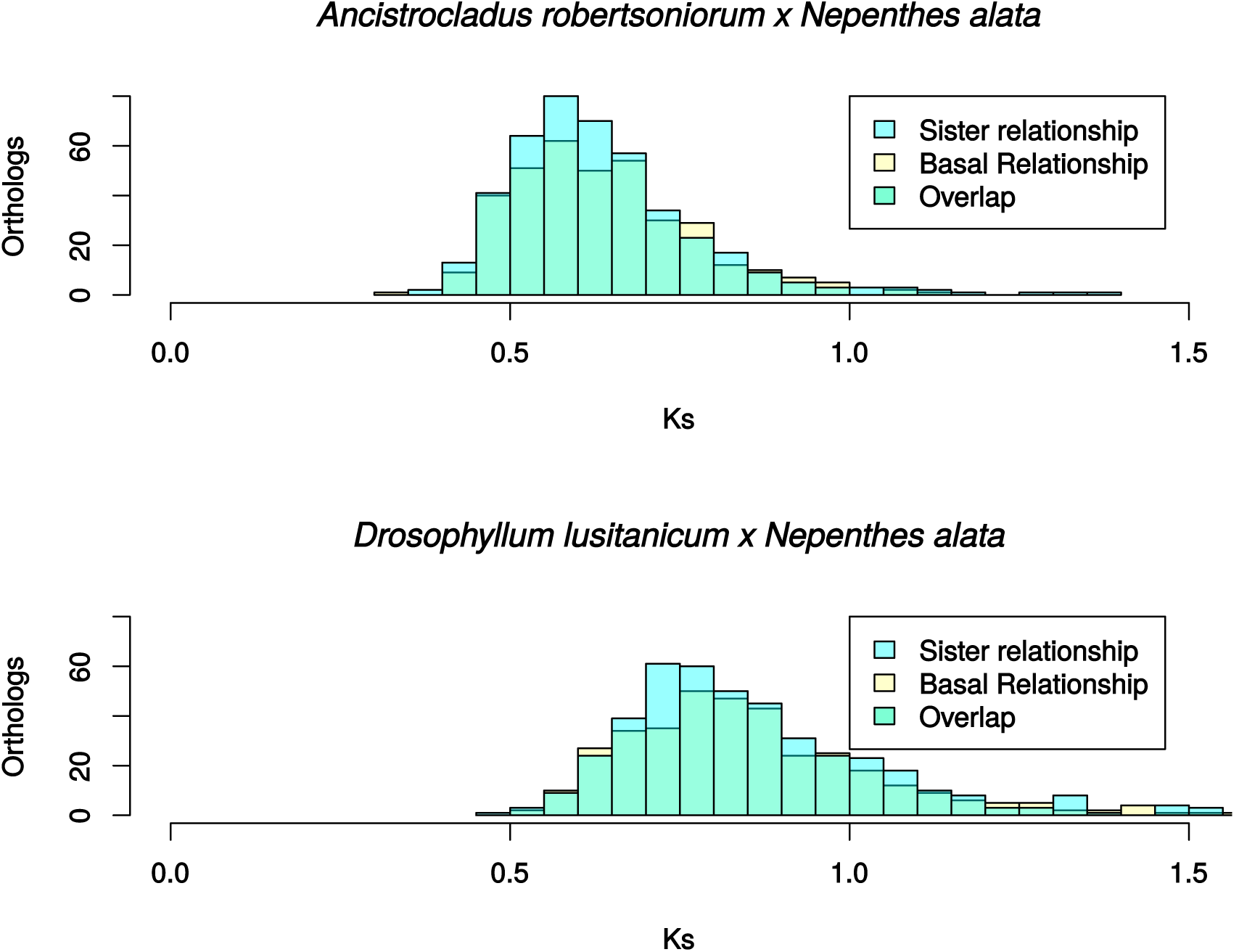
Distribution of synonymous substitutions (Ks values) among conflicting gene tree topologies. Figure shows the distribution of synonymous substitutions between *Nepenthes alata* and *Ancistrocladus robertsoniorum* and the distribution of synonymous substitutions between *Drosophyllum lusitanicum* and *Nepenthes alata.* The values were acquired for the *A. robertsoniorum, D. lusitanicum* and *N. alata* sequences obtained from gene trees that show conflicting topologies of *Drosophyllum* and *Ancistrocladus* sister to *Nepenthes* and *Drosophyllum* and *Ancistrocladus* basal to the rest of the carnivorous Caryophyllales. The mean Ks values for the comparison of *A. robertsoniorum* and *N. alata* were 0.63592 (sister to the other lineages) and 0.6358 (sister to only Nepenthaceae). The mean Ks values for the comparison of *D. lusitanicum* and *N. alata* were 0.85467 (sister to the other lineages) and 0.85861 (sister to Nepenthaceae only).

**Supplementary Figure 4:**
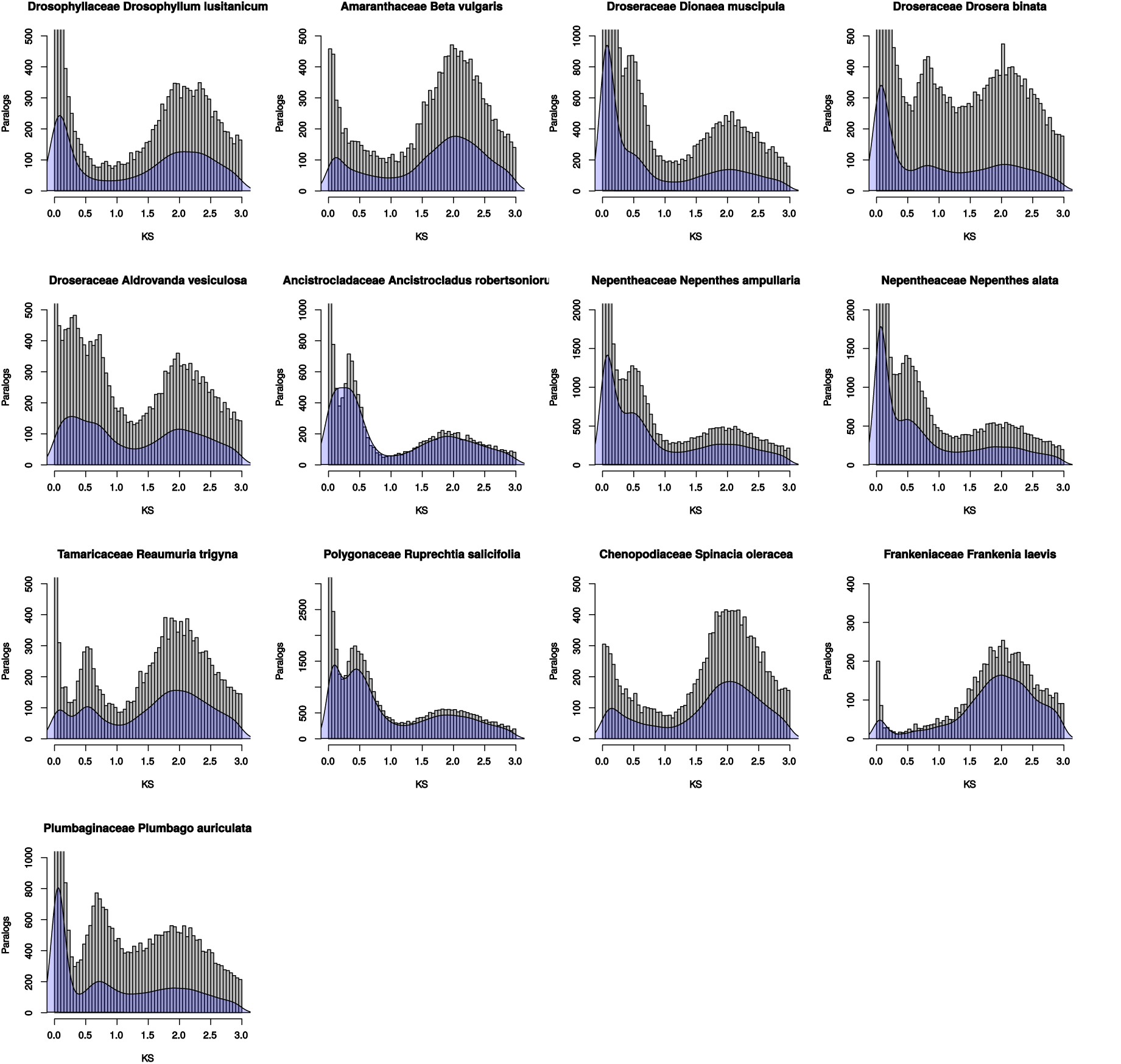
Comparison of synonymous substitutions (Ks values) between inferred paralogs, presented through a histogram (60 bins) with the density plot mapped on top. Comparison of the within species inferred paralogs Ks values as presented in a histogram of 60 breaks and through a superimposed density plot in blue. The Y-axis is for the histograms representing the paralogs with the given Ks value and the Y-axis for the superimposed density plots is not shown. The X-axis represents the Ks value and is the same between the histogram and the density plot.

**Supplementary Figure 5:**
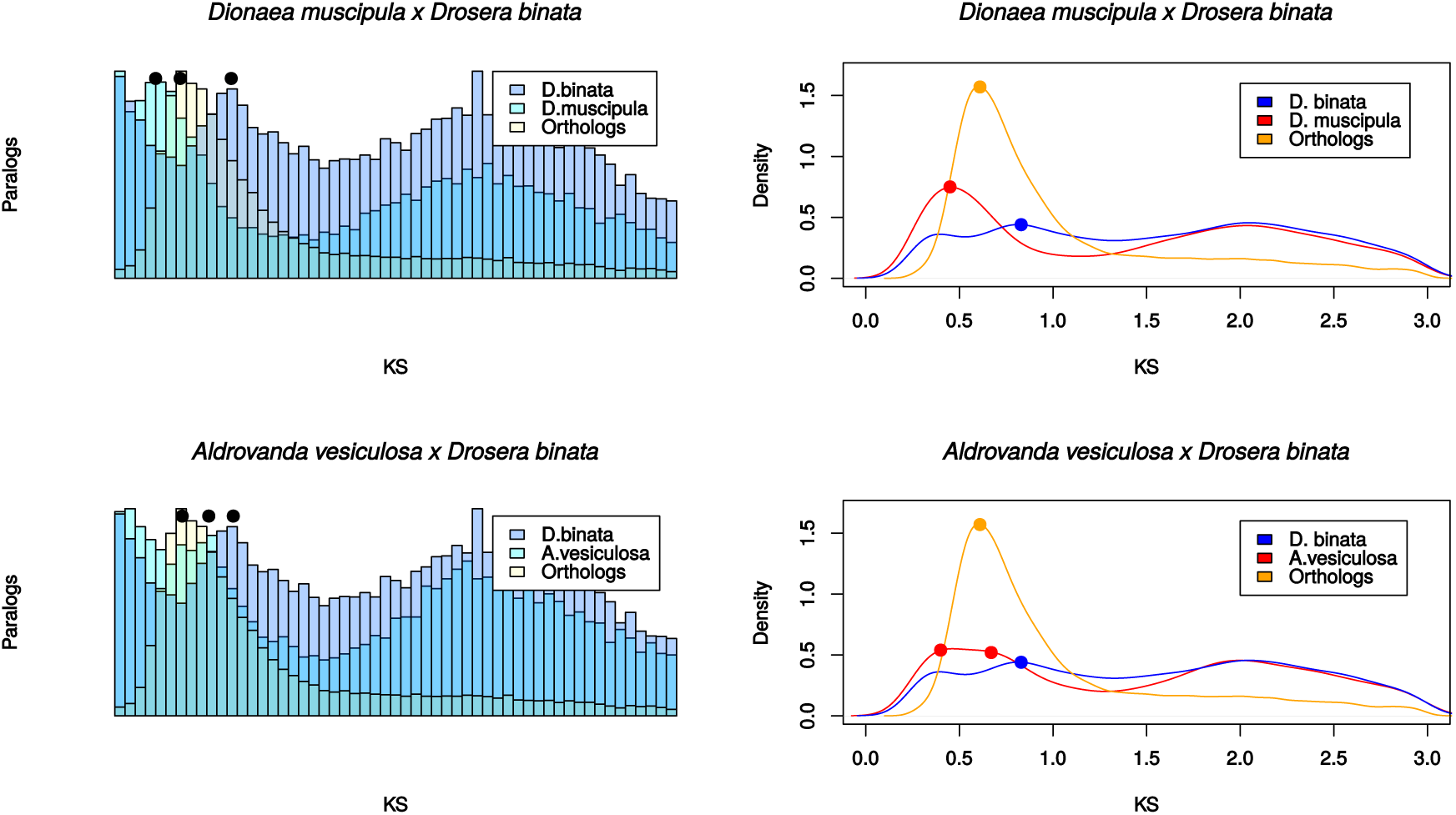
Synonymous substitution (Ks) plots presented as both histogram and density plot for pairwise Droseraceae comparisons. The figure depicts Ks plots between *Drosera binata* and other members of the Droseraceae. Dots are placed on the highest points of the peaks.

**Supplementary Figure 6:**
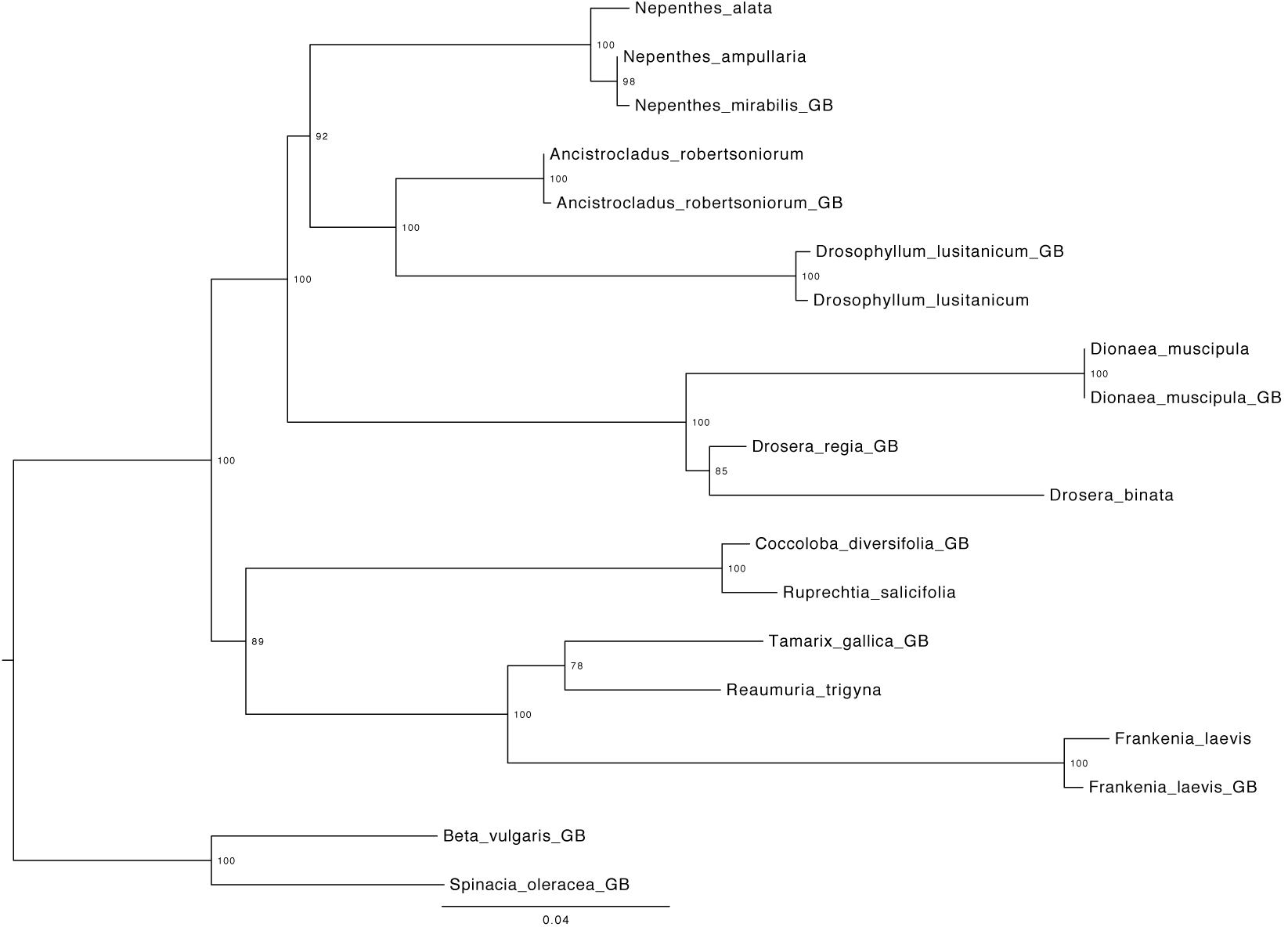
Contamination check of the transcriptomes through the assembly of a maximum likelihood MatK gene tree. The figure shows representative family samples from GenBank (ending in GB) compared the MatK sequence inferred using BLAST from the assembled transcriptome data used in the analyses. The analysis was run for 200 BS replicates with the respective values at the nodes.

**Supplementary Table 1:**
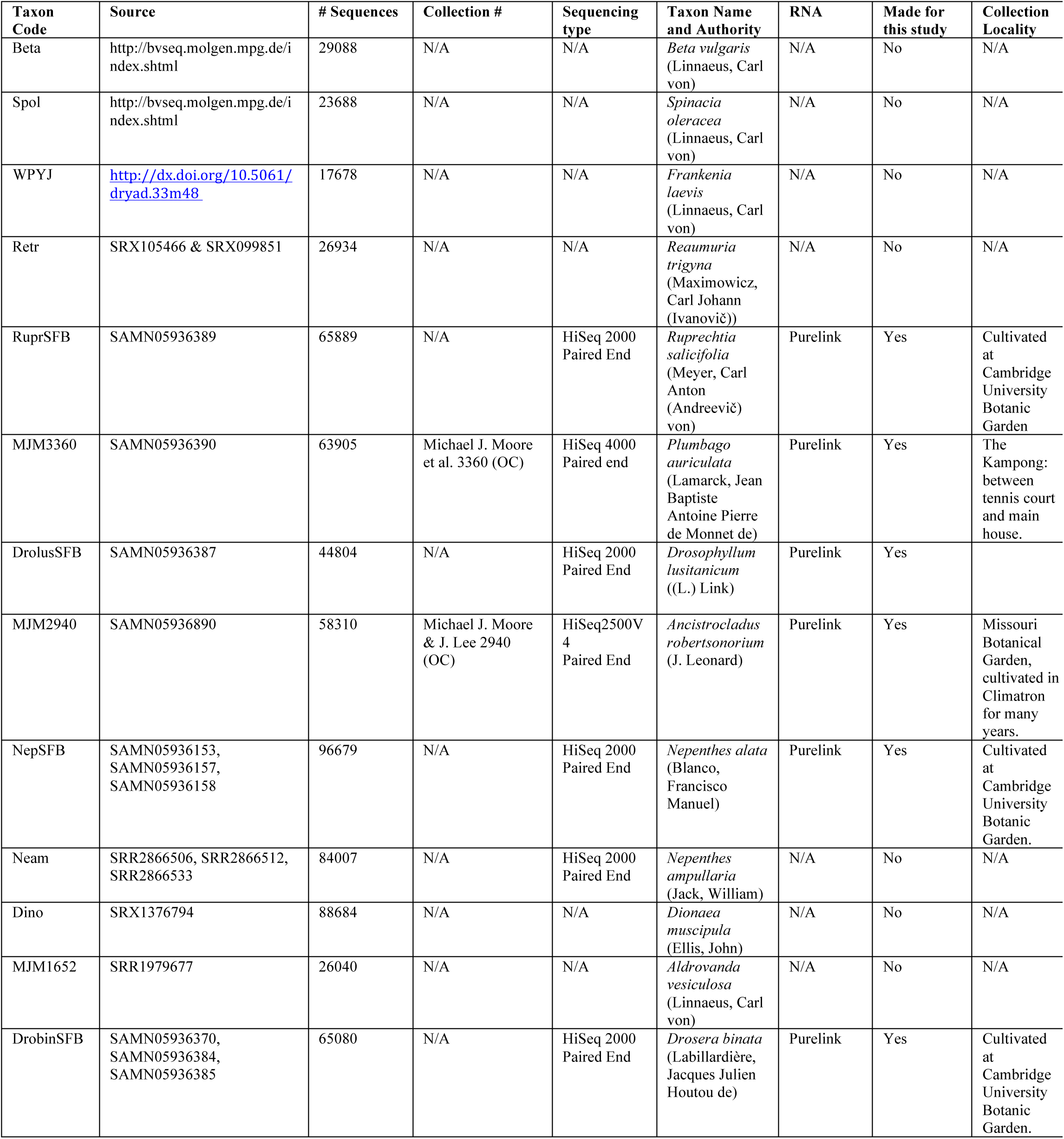
Taxa used for the analyses, sources of data, collections and location of the collections.

| Taxon Code | Source | # Sequences | Collection # | Sequencing type | Taxon Name and Authority | RNA | Made for this study | Collection Locality |
| --- | --- | --- | --- | --- | --- | --- | --- | --- |
| Beta | http://bvseq.molgen.mpg.de/index.shtml | 29088 | N/A | N/A | *Beta vulgaris* (Linnaeus, Carl von) | N/A | No | N/A |
| Spol | http://bvseq.molgen.mpg.de/index.shtml | 23688 | N/A | N/A | *Spinacia oleracea* (Linnaeus, Carl von) | N/A | No | N/A |
| WPYJ | http://dx.doi.org/10.5061/dryad.33m48 | 17678 | N/A | N/A | *Frankenia laevis* (Linnaeus, Carl von) | N/A | No | N/A |
| Retr | SRX105466 & SRX099851 | 26934 | N/A | N/A | *Reaumuria trigyna* (Maximowicz, Carl Johann (Ivanovič)) | N/A | No | N/A |
| RuprSFB | SAMN05936389 | 65889 | N/A | HiSeq 2000 Paired End | *Ruprechtia salicifolia* (Meyer, Carl Anton (Andreevič) von) | Purelink | Yes | Cultivated at Cambridge University Botanic Garden |
| MJM3360 | SAMN05936390 | 63905 | Michael J. Moore et al. 3360 (OC) | HiSeq 4000 Paired end | *Plumbago auriculata* (Lamarck, Jean Baptiste Antoine Pierre de Monnet de) | Purelink | Yes | The Kampong: between tennis court and main house. |
| DrolusSFB | SAMN05936387 | 44804 | N/A | HiSeq 2000 Paired End | *Drosophyllum lusitanicum* ((L.) Link) | Purelink | Yes |  |
| MJM2940 | SAMN05936890 | 58310 | Michael J. Moore & J. Lee 2940 (OC) | HiSeq2500V4 Paired End | *Ancistrocladus robertsoniorum* (J. Leonard) | Purelink | Yes | Missouri Botanical Garden, cultivated in Climatron for many years. |
| NepSFB | SAMN05936153, SAMN05936157, SAMN05936158 | 96679 | N/A | HiSeq 2000 Paired End | *Nepenthes alata* (Blanco, Francisco Manuel) | Purelink | Yes | Cultivated at Cambridge University Botanic Garden. |
| Neam | SRR2866506, SRR2866512, SRR2866533 | 84007 | N/A | HiSeq 2000 Paired End | *Nepenthes ampullaria* (Jack, William) | N/A | No | N/A |
| Dino | SRX1376794 | 88684 | N/A | N/A | *Dionaea muscipula* (Ellis, John) | N/A | No | N/A |
| MJM1652 | SRR1979677 | 26040 | N/A | N/A | *Aldrovanda vesiculosa* (Linnaeus, Carl von) | N/A | No | N/A |
| DrobinSFB | SAMN05936370, SAMN05936384, SAMN05936385 | 65080 | N/A | HiSeq 2000 Paired End | *Drosera binata* (Labillardière, Jacques Julien Houtou de) | Purelink | Yes | Cultivated at Cambridge University Botanic Garden. |

**Supplementary Table 2:** List of species and GenBank accession for the MatK sequences used in the contamination analysis.

| Species | GenBank Accession |
| --- | --- |
| *Drosera regia* | gi\|8568032\|gb\|AF204848.1\| |
| *Dionaea muscipula* | gi\|8568030\|gb\|AF204847.1\| |
| *Nepenthes mirabilis* | gi\|14193614\|gb\|AF315920.1\| |
| *Tamarix gallica* | gi\|8568058\|gb\|AF204861.1\| |
| *Frankenia laevis* | gi\|47498931\|gb\|AY514853.1\| |
| *Coccoloba diversifolia* | gi\|297372635\|emb\|FN597640.1\| |
| *Ancistrocladus robertsoniorum* | gi\|285803889\|gb\|GQ470539.1\| |
| *Drosophyllum lusitanicum* | gi\|47498945\|gb\|AY514860.1\| |
| *Beta vulgaris* | gi\|47498889\|gb\|AY514832.1\| |
| *Spinacia oleracea* | gi\|11497503:1783-3300 |
| *Drosera regia* | gi\|8568032\|gb\|AF204848.1\| |
| *Dionaea muscipula* | gi\|8568030\|gb\|AF204847.1\| |
| *Nepenthes mirabilis* | gi\|14193614\|gb\|AF315920.1\| |

**Supplementary Table 3:**
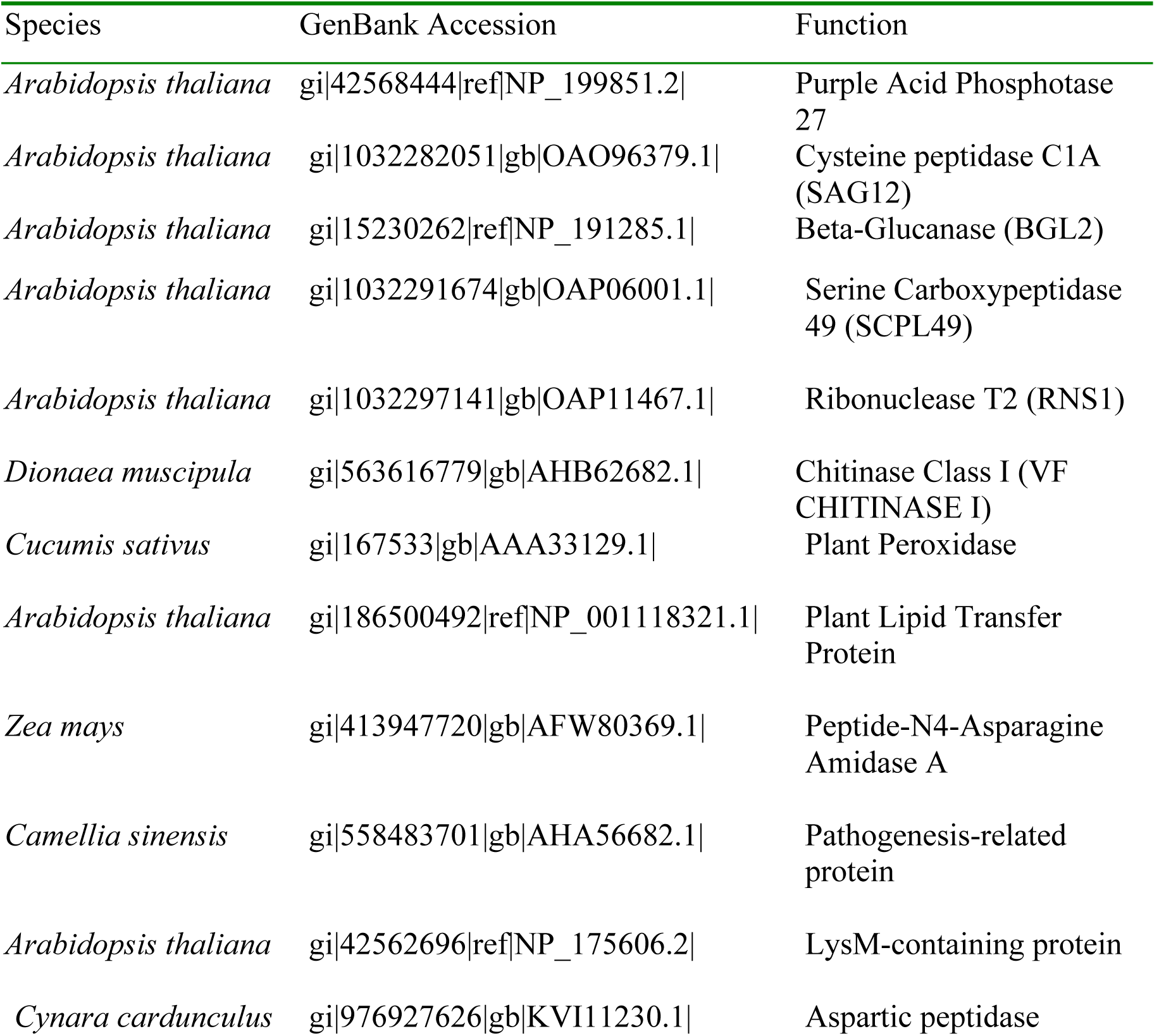
Samples used for identifying homologous clusters of genes identif to be important in carnivory from *Bemm et. al 2016*. Including species name, GenBank acces: and function of sequences.

| Species | GenBank Accession | Function |
| --- | --- | --- |
| *Arabidopsis thaliana* | gi\|42568444\|ref\|NP_199851.2\| | Purple Acid Phosphatase 27 |
| *Arabidopsis thaliana* | gi\|1032282051\|gb\|OAO96379.1\| | Cysteine peptidase C1A (SAG12) |
| *Arabidopsis thaliana* | gi\|15230262\|ref\|NP_191285.1\| | Beta-Glucanase (BGL2) |
| *Arabidopsis thaliana* | gi\|1032291674\|gb\|OAP06001.1\| | Serine Carboxypeptidase 49 (SCPL49) |
| *Arabidopsis thaliana* | gi\|1032297141\|gb\|OAP11467.1\| | Ribonuclease T2 (RNS1) |
| *Dionaea muscipula* | gi\|563616779\|gb\|AHB62682.1\| | Chitinase Class I (VF CHITINASE I) |
| *Cucumis sativus* | gi\|167533\|gb\|AAA33129.1\| | Plant Peroxidase |
| *Arabidopsis thaliana* | gi\|186500492\|ref\|NP_001118321.1\| | Plant Lipid Transfer Protein |
| *Zea mays* | gi\|413947720\|gb\|AFW80369.1\| | Peptide-N4-Asparagine Amidase A |
| *Camellia sinensis* | gi\|558483701\|gb\|AHA56682.1\| | Pathogenesis-related protein |
| *Arabidopsis thaliana* | gi\|42562696\|ref\|NP_175606.2\| | LysM-containing protein |
| *Cynara cardunculus* | gi\|976927626\|gb\|KVI11230.1\| | Aspartic peptidase |

**Supplementary Table 4:**
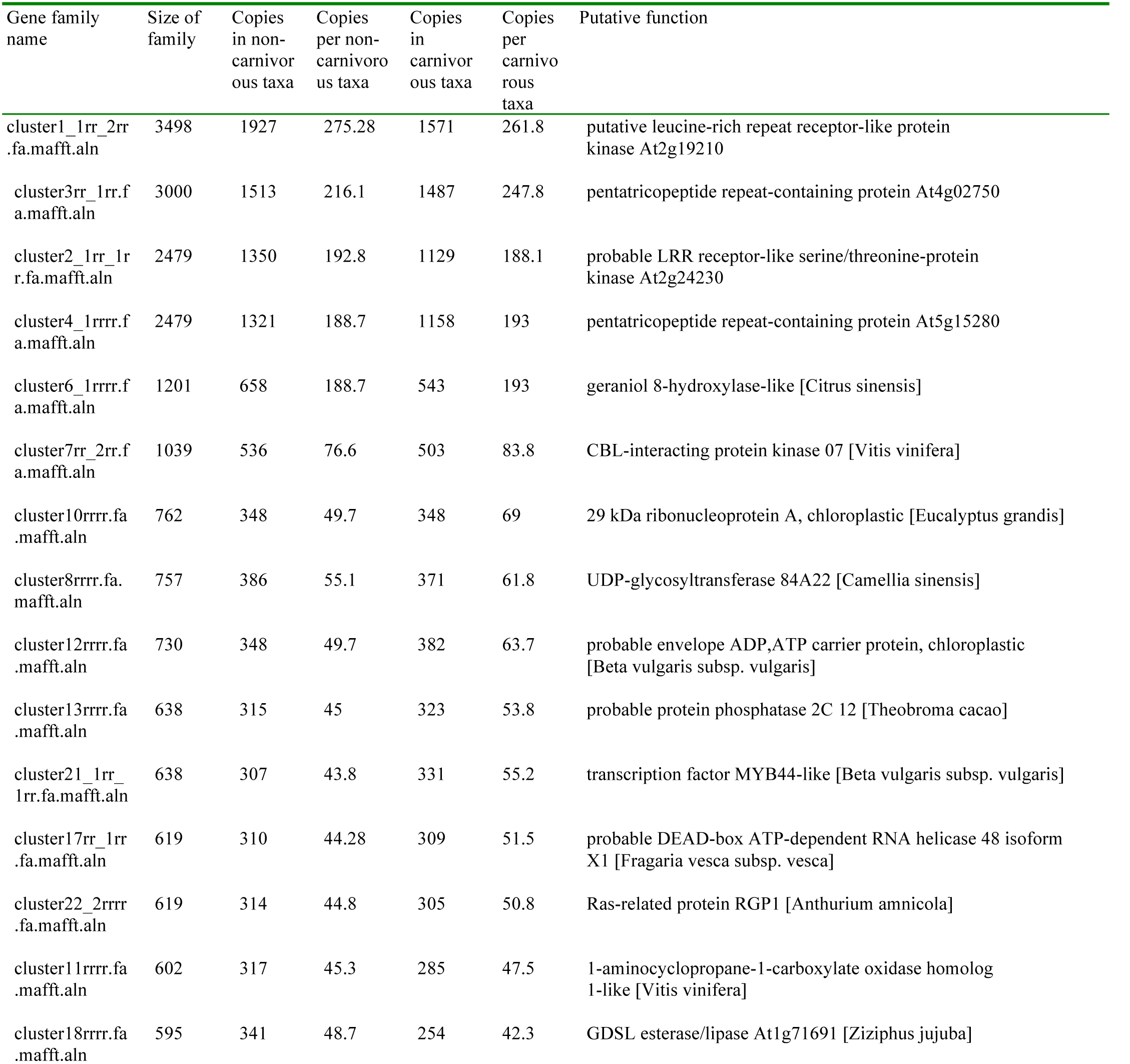
List of largest gene families, divided to size of family found in the carnivorous and non-carnivorous taxa used in the study.

**Supplementary Table 5:** Comparison of gene family size between carnivorous and non-carnivorous taxa identified in carnivory from *Bemm et. al 2016*.

| Name in analysis | Size of family | Copies in non-carnivorous taxa | Average copies per non-carnivorous taxa | Average copies in carnivorous taxa | Average copies per carnivorous taxa | Putative function |
| --- | --- | --- | --- | --- | --- | --- |
| cluster98rrrr.fa.mafft.aln | 199 | 103 | 14.7 | 96 | 16 | Purple Acid Phosphatase 27 |
| cluster82rrrr.fa.mafft.aln | 234 | 113 | 16.1 | 121 | 20.1 | Cysteine peptidase C1A (SAG12) |
| cluster32rrrr.fa.mafft.aln | 416 | 214 | 30.5 | 202 | 33.6 | Beta-Glucanase (BGL2) |
| cluster7000rrrr.f a.mafft.aln | 8 | 3 | 0.4 | 5 | 0.8 | Serine Carboxypeptidase 49 (SCPL49) |
| cluster898rrrr.fa.mafft.aln | 50 | 26 | 3.71 | 24 | 4 | Ribonuclease T2 (RNS1) |
| cluster319_2rrrr .fa.mafft.aln | 62 | 37 | 5.2 | 25 | 4.1 | Chitinase Class I (VF CHITINASE I) |
| cluster24rrrr.fa.mafft.aln | 527 | 324 | 46.2 | 204 | 33.83 | Plant Peroxidase |
| cluster263rrrr.fa.mafft.aln | 108 | 47 | 6.7 | 61 | 10.1 | Plant Lipid Transfer Protein |
| cluster1669rrrr.f a.mafft.aln | 25 | 13 | 1.8 | 12 | 2 | Peptide-N4-Asparagine Amidase A |
| cluster556rrrr.fa.mafft.aln | 69 | 44 | 6.2 | 25 | 4.1 | Pathogenesis-related protein |
| cluster6240rrrr.f a.mafft.aln | 9 | 7 | 1 | 2 | 0.3 | LysM-containing protein |
| cluster439rrrr.fa.mafft.aln | 70 | 28 | 4 | 42 | 7 | Aspartic peptidase |

**Supplementary Table 6:**
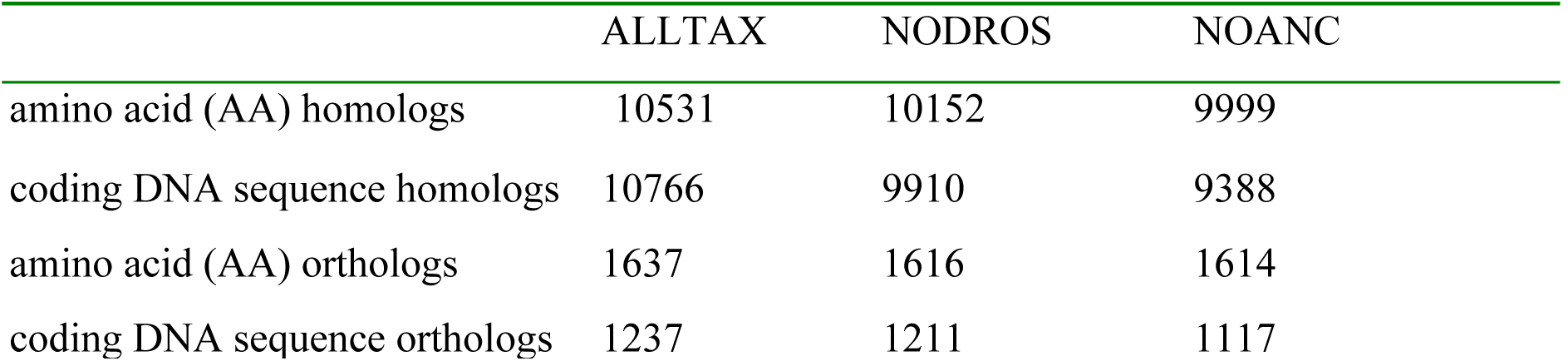
Composition of datasets used for the phylogenomic analyses.

|  | ALLTAX | NODROS | NOANC |
| --- | --- | --- | --- |
| amino acid (AA) homologs | 10531 | 10152 | 9999 |
| coding DNA sequence homologs | 10766 | 9910 | 9388 |
| amino acid (AA) orthologs | 1637 | 1616 | 1614 |
| coding DNA sequence orthologs | 1237 | 1211 | 1117 |

